# An optimised method for generation of murine CAR-T cells by CRISPR/Cas9

**DOI:** 10.1101/2025.11.29.689298

**Authors:** Thomas J Jackson, Courtney Himsworth, Sophie Munning-Tomes, Farah Alam, Helena Brezovjakova, Laura K Donovan, Amy K. Erbe, Paul M. Sondel, Louis Chesler, John Anderson

## Abstract

Development of the next generation of chimeric antigen receptor (CAR) T-cells requires assessment in systems that better recapitulate the suppressive tumour microenvironment of solid tumours. CRISPR-Cas9 knock-in of promoter-less homology directed repair templates (HDRT) into the T-cell receptor locus has been shown to result in physiological expression of CARs with improved tumour control. We initially compared the use of dsDNA and adenovirus associated virus (AAV) HDRTs in mouse T cells. We have subsequently developed an optimised method for AAV transduction resulting in high editing efficiencies with minimal toxicity. In contrast with our experience of retroviral transduction of mouse T cells, our CRISPR/Cas9 AAV transduction method results in sustained CAR expression and T cell expansion in vitro as well as in vivo persistence. This approach allows for pre-clinical assessment of individual and libraries of CAR constructs in relevant immune-competent mouse models.

## Introduction

CAR-T cell therapy has revolutionised treatment of relapsed/refractory B cell acute lymphoblastic leukaemia^1,2^. Whilst there have been some notable successes in some paediatric solid tumours such as neuroblastoma^3,4^, there are significant remaining challenges for solid tumour CAR-Ts including tumour homing, the immune suppressive TME, T cell exhaustion^5^.

To develop and test the next generation of CARs that are armoured to overcome these challenges requires model systems that closely model the TME. Experimental approaches to reproduce the TME stress conditions of repeated antigenic stimulation of CAR-T cells in vivo, using repeat stimulations assays in vitro, lack many of the suppressive cells, accessibility barriers and nutrient deficiencies of relevant tumour tissues. Consequently, data from such approaches correlates poorly with in vivo function^6^, further underscoring need for in vivo immune competent models.

CRISPR-Cas knock-in uses a ribonuclear protein (Cas9 and sgRNA guide) to generate double strand break (DSB) at a specific locus, which in the presence of a repair template results in gene knock in by homology-directed repair (HDR). This HDR-based approach has at least two advantages. First, a CAR can be placed under the control of an endogenous promoter with favourable characteristics. For example, expression of CARs placed in the TRAC locus are down regulated upon activation, reducing exhaustion and increasing tumour efficacy^7^. Second, this technique readily supports library-based approaches due to the guaranteed single-site integration, whereas retroviral based approaches require limiting dilutions to ensure single integration per cell resulting in low transduction rates and variability of transcriptional regulation. Library screens have been used to identify CAR designs in human T cells with increased persistence in immune-deficient models^8,9^.

Homology directed repair templates have been delivered to the cell as electroporated dsDNA^10^ or as an adeno associated virus (AAV) vector^7^. For dsDNA templates, efficiencies can be improved by addition of truncated cas9 target sequences (tCTS), which can be bound by RNP without inducing cleavage, presumably by enhanced nuclear localisation of the HDRT^11^. By contrast, AAV virions are internalised by endocytosis due to cell surface receptor-capsid protein interactions, undergo endosomal escape and nuclear localisation prior to uncoating and genome release^12^. There have been reports that the internal tandem repeats (ITRs) of AAV HDRT can result in concatenation prior to integration and that addition of a complete CTS (i.e. cleavable) adjacent to the ITRs mitigates this^13^.

HDR occurs in late S and G2 phase and therefore gene editing utilising this pathway is more efficient in dividing cells, such as activated T cells. However, following T cell activation there is increased activity of the cytosolic DNA sensing cGAS-STING pathway driving cell death or senescence. It has therefore been suggested there is a goldilocks-zone for gene editing of primary activated T cells using HDRT, which lies after activation but prior to full stress response^14^. HDR can be favoured by inhibition of alternative double strand break repair pathways such as non-homologous end joining and microhomology mediated end joining^15^.

In this paper we systematically compare dsDNA vs AAV HDRT for editing of mouse T cells and then describe the development of an optimised method for generating murine TRAC-CAR T cells for use in immunocompetent mouse models.

## Methods

### Animals

C57Bl6NTac (MGI:2164831), 129SvJ/X1 (MGI:3044210) and C57Bl6J-Rosa26eGFPCas9 (JAX #:026179) mice colonies were bred and maintained from existing stock. FVBNrj were purchased from Janvier Labs. All animals were housed in individually ventilated cages with water and food available *ad libitum* and monitored daily. All procedures were reviewed and approved by a University College London (UCL) biological services named animal care and welfare officer and named veterinary surgeon and performed in accordance with Home Office project license PP5675666.

### Generation of human B7H3 expressing murine neuroblastoma

Human B7H3 transgenic mice (C57Bl6NTac-hB7H3^fl/fl^) were obtained from Taconic Biosciences. Briefly, mouse embryonic stem cells were edited by homologous recombination into the murine B7H3 locus with a vector containing a chimeric B7H3 exon 2 with the mouse 5’ untranslated region (UTR) and human coding DNA sequence (CDS) followed by lox-STOP-Lox, an intron-less human B7H3 CDS, mouse 3’ UTR and a polyadenylation (pA) signal. These mice were crossed with 129SvJ-Th-MycN, Th-Cre mice to generate mice with neural crest lineage restricted human B7H3 and MycN expression. A spontaneous tumour from a C57Bl6J;129SvJ(N2)-Th-MycN^+/-,^ Th-Cre^+/-^ hB7H3 ^fl/wt^ mouse was taken and disaggregated using Accumax (ThermoFisher) and in vitro passaged to create the tumoursphere line 370566.

### Cell culture

All cells were cultured in sterile tissue-culture treated plates and flasks, in an incubator at 37°C 5% CO_2_.

AAVPro (Takara) were cultured in DMEM 4.5g/L glucose supplemented with 10% fetal calf serum (FCS), 2mM GlutaMAX, and penicillin/streptomycin. Phoenix Eco were cultured in IMDM with 10% FCS 2mM glutamine.

9464D were cultured in DMEM 4.5g/L glucose supplemented with 10% FCS, 2mM glutamine, 1X non-essential amino acids (Sigma), 1mM sodium pyruvate. Two derivative cell lines expressing either humanB7H3 and mCherry (9464D-B7H3-mCherry) or GD2 and GD3 synthases (9464D-GD2) were maintained in the presence of 6μg/ml puromycin, or 6μg/ml puromycin and 7.5μg/ml blasticidin respectively^16^.

370566 tumourspheres were cultured in DMEM/F12 supplemented with 15% FCS,1X penicillin/streptomycin, 1X B27 supplement (ThermoFisher), 50μM β-mercaptoethanol, 15ng/ml human b-FGF and 10ng/ml human EGF (Preprotech) (adapted from Cao et al. ^17^). Spheroids were passaged by disaggregation using accutase (Sigma).

Mouse splenocytes were isolated by passing spleens through a 70μm cell strainer with the back of a sterile 1ml syringe plunger. Strainers were washed with PBS supplemented with 5mM EDTA and splenocytes pelleted and resuspended in ACK lysis buffer and incubated at room temperature for 5 minutes. Cells were pelleted and resuspended in FACS buffer (2% FCS+ 1mM EDTA in PBS), passed through a further 70μm cell strainer to obtain a single cell suspension. T cells were then isolated by negative selection using the Mojosort mouse T cell isolation kit (Biolegend). Mouse T cells were cultured in RPMI supplemented with 10% FCS, 1X penicillin/streptomycin, 2mM GlutaMax, 50μM β-mercaptoethanol. T cell activation was performed either by addition of anti-CD3 anti-CD28 dynabeads (ThermoFisher) as per the manufacturer’s instructions or with purified antibodies in a 24 well format. For antibody activation 0.5ml/well of 1μg/ml anti-CD3 (Ultra-LEAF clone 145-2C11, Biolegend) was applied for 2 hours at 37°C and then washed prior to application of cells in media supplemented with 5μg/ml anti-CD28 (Ultra-LEAF Clone 37.51; Biolegend). During T cell activation mouse T cell media was supplemented with 50U/ml human IL-2. During T-cell transduction and expansion media was supplemented with 50U/ml human IL-2, 5ng/ml mouse IL7 and 10ng/ml human IL15 (PreProtech)^6^. During expansion T cell density was maintained at ∼5x10^5^cells/cm^2^ and 1-2x10^6^cells/ml.

### Cloning

A 989bp region of the murine TRAC locus (NCBI Reference Sequence: NG_007044.1 position 1792597 to 179358) was obtained as a geneblock (Twist biosciences). This was cloned into a pUC19 vector using HiFi cloning (NEB). This was further subcloned to generate pUC19 vectors with 300bp homology arms and promoter-less transgenes inserted at the cut site in TRAC exon 1 of the sgRNA guide with sequence 5’-AGGGTGCTGTCCTGAGACCG-3’. Transgenes included: a) eGFP; b) a membrane bound mini-luciferase formed by fusion of teLuc^18^ and the transmembrane domain of mouse CD28 with a CD28 signal peptide; c) CARs with scFvs targeting GD2 (huk666)^19^ or human B7H3 (TE9)^20^ with a murine CD8 stalk, CD28 and CD3z endodomains, with or without a C-terminal degron (iTag2). CAR constructs were co-expressed with either the suicide switch RQR8^21^ or RMR8, a homolog with the QBend10 epitope replaced with a Myc-Tag (EQKLISEEDL). To generate AAV transfer plasmids, required regions of pUC19 based vectors were amplified using Q5 polymerase (NEB) with primers containing MluI and RsrII sites and cloned into the pAAV-minCMV-mCherry backbone (a gift from Feng Zhang (Addgene plasmid # 27970)). For vectors including an sgRNA expression cassette, geneblocks with either U6 or M11 promoters^22^ upstream of the Cas9 sgRNA scaffold, TRAC guide and polyA signal were used. pAdDeltaF6 was a gift from James M. Wilson (Addgene plasmid # 112867). Ark313 was a gift from Justin Eyquem^23^ (Addgene plasmid # 200257). pMiniHelper was generated by HiFi cloning from 5 fragments of pAdDeltaF6. AAV transfer plasmids were propagated in NEBStable, all other plasmids were in NEB5α. Plasmids sequences were confirmed by whole plasmid sequencing (Plasmidsaurus).

### dsDNA HDRT generation

dsDNA HDRT were generated by Q5 PCR with primers targeting the homology arms and containing tCTS in an in-in orientation but without edge sequences^11^. DNA was purified using nucleospin columns (Machery Nagel), then concentrated by sodium acetate/isopropanol precipitation and resuspended in hybridisation buffer (IDT) to a concentration of 1μg/μl.

### AAV production and titre

For smaller scale generation of ∼40ul concentrated (>1x10^13^ Vg/ml) AAV preparations the protocol of Regrini et al.,^24^ was used with modifications. AAV Pro cells were cultured in two 175cm^2^ flasks and transfected when 70% confluent using PEIStar (Tocris) with 9.2fmol/cm^2^ of each of the RepCap plasmid Ark313, a helper plasmid (pAdDeltaF6 or miniHelper) and AAV transfer plasmid. After 24 hours 90% of the media was replaced with OptiPro SFM (ThermoFisher). 72 hours post transfection AAV was harvested from media and cells using PEG-chloroform isolation. Following resuspension in 50mM HEPES pH 8 the crude AAV extract was treated with 50U/ml benzonase nuclease with 2.5mM MgCl_2_ at 37°C for 30 minutes. AAV was isolated by 4 rounds of chloroform extraction followed by buffer exchange into PBS supplemented with 2.5mM KCl, 1mM MgCl_2_ and 0.001% F68 pluronic acid (PBS-MK-F68) using 2ml 100kDa centrifugal filters (Amicon).

For larger scale preparations transfection was performed as above but using twenty 15cm dishes. 72 hours post transfection, cells were pelleted and resuspended in 5ml lysis buffer (140mM NaCl, 5mM KCl, 3.5mM MgCl2, 25mM Tris pH 7.5) and lysed by 3 freeze thaw cycles. AAV was precipitated from supernatants by adding 31.3g/100ml ammonium sulphate and incubating at 4°C for 30 minutes then centrifuged at 5200xg 45 min 4°C and resuspended to a final volume of 15ml with lysis buffer. Both crude extracts were combined and treated with 50U/ml benzonase nuclease at 37°C for 1 hour. The treated extracts were applied to discontinuous (10%-25%-40%-60%) iodixanol gradients in six 13.2ml (14x89mm) ultracentrifuge tubes (Beckman Coulter) and centrifuged in a TH-641 swinging bucket rotor at 41000rpm for 2 hours 20mins at 18°C. The tubes were pierced to extract the 40% layer, avoiding the 25-40% interphase and buffer exchange to PBS-MK-F68 performed using 15ml 100kDa centrifugal filters (Amicon) in a final volume of 0.5-1ml.

AAV preparations were diluted 1:1,000, 1:10,000 and 1:100,000 in water then quantified by qPCR using SsoAdvanced Universal SYBR Green Supermix (biorad) and primers for the ITR (5’-GGAACCCCTAGTGATGGAGTT-3’ and 5’-CGGCCTCAGTGAGCGA-3’). Linearised transfer plasmid was used to generate a standard curve of 10^9^ – 10^2^ molecules/μl. AAV preparations were stored in 25-50μl aliquots at -80°C and could be used for up to 2 freeze thaw-cycles.

### Retrovirus production and transduction

Pheonix Eco cells were transfected at 70-80% confluency using GeneJuice (sigma) with a SFG transfer plasmid containing a CAR with a B7H3 scfv (376.96)^25^ mouse CD8 stalk, CD8 transmembrane and CD28 and CD3z endodomains (“376.96m28z”). Supernatants were harvested at 48- and 72-hours post transduction and pooled. Splenocytes were activated for 48 hours with anti-CD3/anti-CD28 Dynabeads as per manufactured instructions then transduced with 1.5ml virus/1.5x10^6^ cells in rectronectin coated plates with spinoculation at 1000xg for 40 minutes.

### Electroporation

Splenic T cells were activated for 48 hours with anti-CD3/anti-CD28 Dynabeads as per the manufacturer’s instructions then electroporated using the P4 Primary Cell 4D-nucleofector Kit S (Lonza). Briefly, following removal of Dynabeads, cells were pelleted at 100xg for 10mins room temperature and resuspended at 1.25x10^6^ cells/20μl P4 buffer (Lonza). 100pmol each of TRAC exon 1 sgRNA (5’-AGGGUGCUGUCCUGAGACCG-3’) (IDT) and Alt-R Sp HiFi Cas9 (IDT) were combined with PBS to a final volume of 5μl and left for 15 minutes to form RNP complexes. 20μl of cell suspension was added to 5μl RNP, mixed by gentle pipetting then 20μl electroporated using code CM137. Post electroporation cells were recovered by addition of 100μl of T cell expansion media to the cuvette wells and incubated at 37°C for 20 minutes, then transferred to a 48 well plate in a final volume of 500μl with or without 1μM AZD7468 and/or 3μM PolQi2 (MedChemExpress).

For homology directed repair, dsDNA HDRT was added to a final concentration of 5-80nM prior to electroporation or else AAV was added at an MOI of 10^3^-10^5^ Vg/cell after electroporation recovery. Media was changed 24 hours after electroporation.

### Electroporation free AAV transduction

For electroporation free AAV transduction, activated splenic T cells from C57Bl6-Rosa26eGFPCas9 mice were transduced with AAVs containing a HDRT and TRAC sgRNA under either a U6 or M11 promoter with or without AZD7468/PolQi2. Media was changed after 24 hours.

### Flow cytometry

Flow cytometry was performed on the Cytoflex V4-B2-Y4-R3 Flow Cytometer (Beckman Coulter). Cells were stained in FACS buffer (PBS 2% FCS 1mM EDTA) using the antibodies in Supplementary table 1. Either DAPI or Ghost Red780 (Tonbio) were used as viability dyes. A minimum of 5000 live events were analysed for in vitro assays. 20μl EDTA treated whole blood was processed for *in vivo* studies and all events acquired.

### Co-coculture assays

9464D-GD2 or 9464D-mCherry hB7H3 were plated at 25000/well in 96 well flat bottom plate in 9464D media without selective antibiotics. 2 hours later, 100μl of CAR-T in RPMI 10% FCS 2mM Glutamax were added at 1:1 or 2:1 effector: target and cultured for 24-48 hours. T cells were removed, wells washed with PBS and target cells detached by application of trypsin for 5 mins at 37°C. Target cell numbers were quantified by flow cytometry of a fixed volume. Cell killing was calculated as percentage of live cells relative to the mean cell number in no effector wells.

### In vivo assessments

370566 tumorspheres were disaggregated and dead cells removed by overlaying the cell suspension onto ficol and centrifugation at 1400xg for 20 minutes with no brakes. The live cell layer was removed and resuspended at 5x10^6^ cells/ml in ice cold 50% Matrigel LDEV-free (Corning) 50% PBS and kept on ice prior to implantation. 100μl cell mix was injected subcutaneously in the shaved flank of female 129SvJ:C57Bl6J-eGFPCas9 F1 hybrid mice under anaesthesia. 21 days after engraftment, mice with palpable tumours (approximately 50-100mm^3^) were lymphodepleted by intraperitoneal injection of 150mg/kg cyclophosphamide in PBS^26^. Mice were randomised by Latin square based on tumour volume to either no further treatment, 5x10^6^ untransduced T cells or 5x10^6^ CAR-T cells, administered by tail vein injection 24 hours after lymphodepletion. Tumour volumes (width^2^x length/2) were monitored by calliper measurements 3 times/week. Blood was taken by saphenous vein bleeds at 3, 10 and 17 days after CAR-T administration and assessed for CAR persistence by flow cytometry.

### Statistical analysis

Data was analysed with PRISM or in the R statistical environment using a custom script. Where appropriate, differences were assessed using one-way and two-way ANOVAs.

## Results

### AAV is superior to dsDNA as a HDRT in activated mouse T cells

To compare the efficiency and toxicity of dsDNA and AAV as HDR templates mouse splenic T cells were isolated, activated for 48 hours, electroporated with a Cas9 RNP targeting exon 1 of the T cell receptor constant (TRAC) locus and a promoter-less GFP HDRT was delivered as either dsDNA during electroporation or AAV post electroporation (Figure 1A). Flow cytometry 4 days following electroporation demonstrated loss of CD3 and gain of GFP expression in HDRT treated cells (Figure 1B). We found a dose dependent negative effect of dsDNA and the presence of NHEJ and MMEJ inhibitors on cell viability (Figure 1C). Only at 80nM dsDNA were we able to detect low levels of GFP knock-in (1.5% without inhibitors and 3.5% with both inhibitors) but with poor viability (Figure 1C and 1D). By contrast AAV transduction had little effect on cell viability, but up to 80% knock-in rates could be achieved in the presence of both inhibitors.

Previous work with the modified AAV capsid Ark313 had implicated Qa2 expression as crucial for AAV transduction^23^. We therefore assessed the Qa2 expression in 3 murine strains used for immune competent modelling – C57Bl6, 129SvJ and FVBNrj. All expressed cell surface Qa2, but FVBNrj expressed lower levels, which correlated with lower knock-in rates (Supplementary Figure 1).

**Figure 1.**
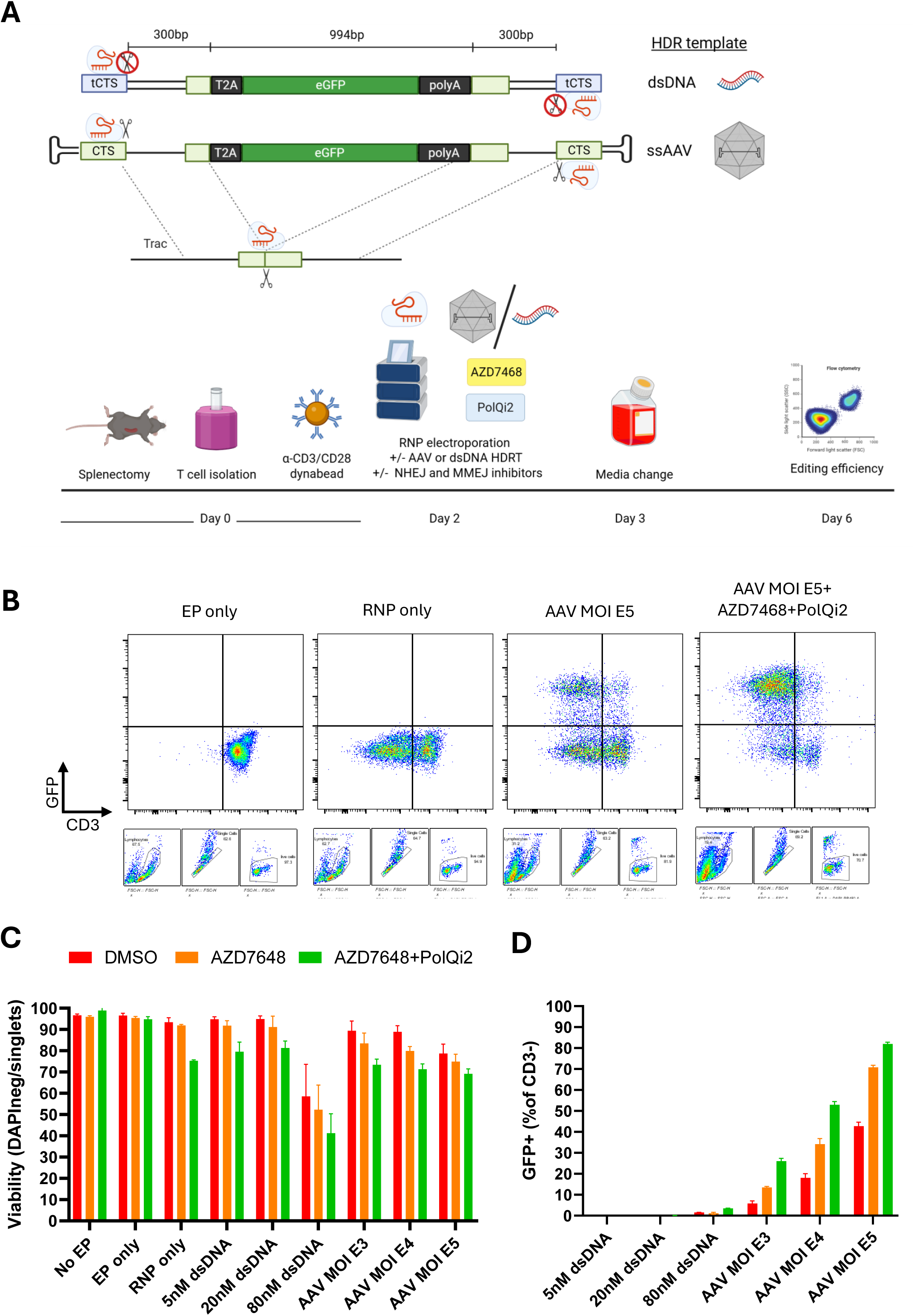
A) Mouse splenic T cells were isolated, activated for 48 hours then electroporated with a Cas9 RNP targeting TRAC +/- promoter-less homology directed repair templates (HDRT) +/- inhibitors of alternative double strand break repair mechanisms. HDRTs were either dsDNA with truncated cas9 target sites (tCTS) to aid localisation or AAVs with full CTS to reduce vector concatenation. Inhibitors were removed after 24 hours by media change and editing efficiency determined by flow cytometry. Spleens from up to 5 animals of the same strain were used (3 for C57Bl6TacN 3 for FVB and 5 for 129SvJ). Data in this figure are for C57Bl6TacN B) Representative flow cytometry plots, gated on single live cells. C) Cell viability by flow cytometry, gated on single cells. Data are means of technical replicates. N = 2 for dsDNA and n=3 for AAV D) Knock-in efficiency normalised to knock-out efficiency (GFP+ cells as percentage of CD3-).

### A miniaturised high copy number AAV helper plasmid supports high titre viral production

Having established the superiority of AAV as a HDRT in murine T cells, we next looked to optimise the generation of AAV. AAV production requires the co-expression of the adenoviral genes E1A, E2A, E4 ORF-6, E4 ORF-6/7, L4-22kDa and VA RNA_I_. HEK293T cells express E1A and the remainder of the genes are delivered by transfection of a helper plasmid. The commonly used helper plasmid pAdDeltaF6 is low copy number and contains extraneous portions of the adenoviral genome. We generated a miniature helper plasmid designated MiniHelper through condensation of essential components into a pUC19 backbone to yield a high copy number vector. When used in AAV production MiniHelper yielded similar viral titre to that obtained with the pAdDeltaF6 (Supplementary Figure 2).

### Antibody based T cell activation results in higher editing efficiencies than CD3/CD28 dynabead activation

We hypothesised that greater viability and yield of mouse CAR-T cells could be achieved using electroporation-free transduction. To that end, activated T cells from transgenic mice that constitutively express cas9 and GFP were transduced with AAVs containing a bicistronic GFP-nano-luciferase HDRT and the sgRNA under either a U6 or M11 promoter (Figure 2A). GFP knock-in was significantly lower from AAVs with the sgRNA under the M11 promoter (Figure 2B) whilst assessment of mini-luciferase activity was limited by solubility of the substrate. Knock-in efficiencies were higher after 36 or 48 hours of activation compared to 24 hours, and activation with plate-bound anti-CD3 and soluble anti-CD28 yielded higher knock in efficiency compared with CD3/CD28 Dynabeads (Figure 2C).

**Figure 2.**
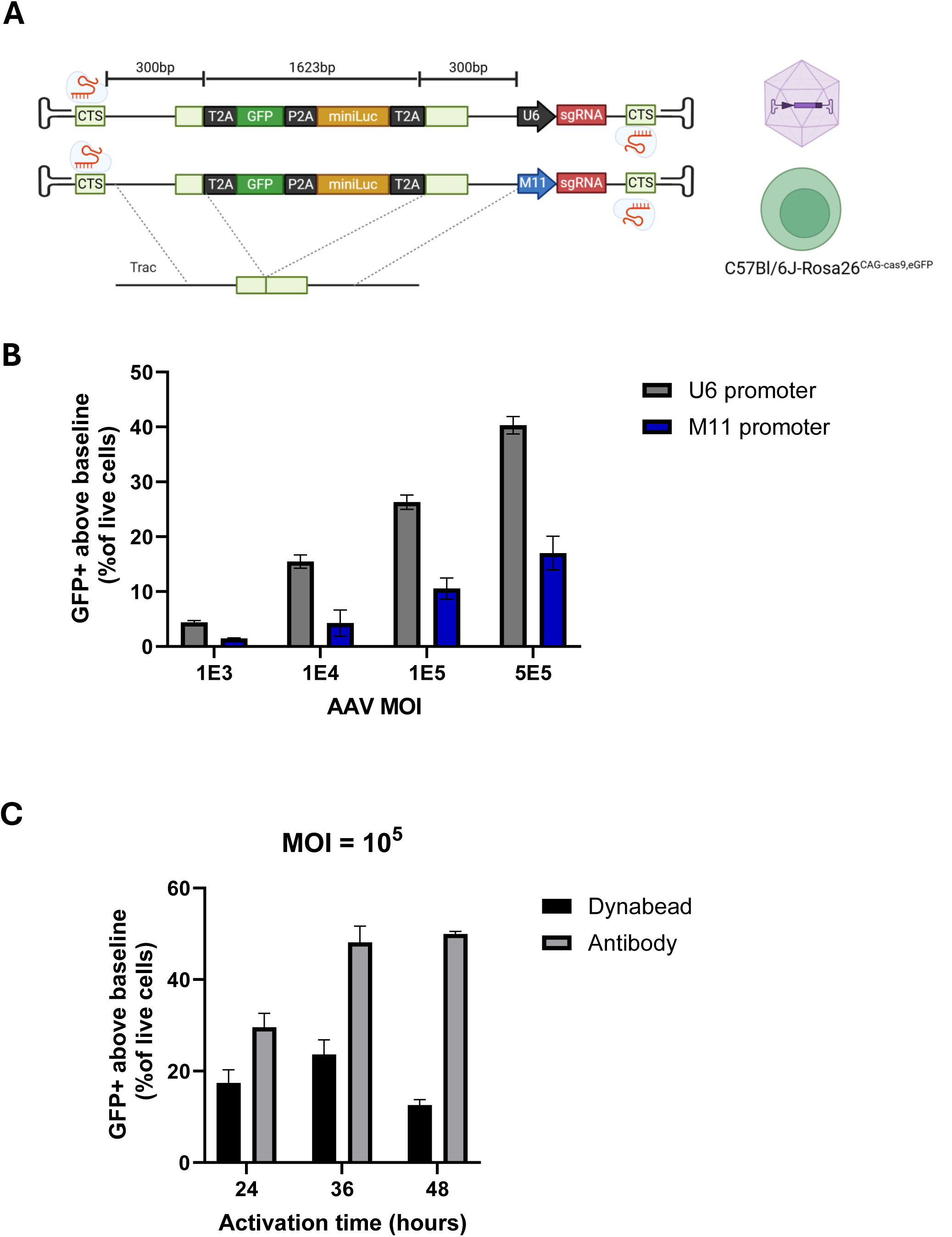
A) AAV HDRT template with polycistronic construct (GFP and miniature membrane bound Luciferase) and sgRNA under either U6 or M11 (H1-7SK hybrid) promoters B) Splenic T cells from transgenic cas9 mice were activated with dynabeads and transduced with the specified MOI of AAV after 48 hours in 1uM AZD7468 and 3uM PolQi2. Percentage of cells expressing GFP above baseline level were determined by flow cytometry 5 days after transduction. Data are mean of 3 technical replicates with standard errors. C) Cas9 mouse T cells were activated with either CD3/CD28 dynabeads or platebound anti-CD3 and soluble anti-CD28 transduced at an MOI 1E5 with a U6 containing AAV at either 24 36 or 48 hours of activation. GFP positivity was assessed by flow cytometry 5 days post transduction. Data are mean of 3 technical replicates with standard errors

### CRISPR/Cas9 mediated knock in results in sustained CAR expression, effective *in vitro* cytotoxicity and the ability to tolerate cryopreservation

Using our optimised activation protocol and AAV transfer plasmid design, we assessed the editing efficiency for HDRTs for CARs targeting either GD2 or human B7H3. We tested a range of different constructs, including those with a degron currently being developed for a Phase I trial (iTag2, a derivative of the published iTAG1 sequence^27^), and compared constructs with and without flanking CTS (Figure 3A). Of note, we identified a population of cells following transduction with all CAR constructs that were CAR-positive but CD3 positive, just as we had with the GFP based HDRT. However, we note that this population is negative for TCRβ (Figure 3B), and therefore we infer that these are likely γδT cells as has been previously shown for TRAC-CARs^28^. All constructs resulted in high (∼90%) transduction efficiencies which were sustained for 15 days *in vitro* (Figure 3C). By contrast, expression of a murinised B7H3 CAR construct delivered by retroviral transduction was not sustained beyond day 7 (Supplementary Figure 3A-C). This loss of CAR expression from retroviral constructs was not due to transient pseudo-transduction as no expression was observed in the presence of retroviral protease and integrase inhibitors (Supplementary Figure 3D). AAV transduced cells expanded approximately 10-fold by day 6, similar to untransduced activated T cells (Figure 3D). Both GD2 and B7H3 CARs showed antigen restricted cytotoxicity against a murine neuroblastoma cell line expressing either GD2 or human B7H3 (Figure 3E). Furthermore, we were able to demonstrate that following cryopreservation of these murine CAR-T cells, they retained viability and cytotoxic potential post thawing (Figure 3F)

**Figure 3.**
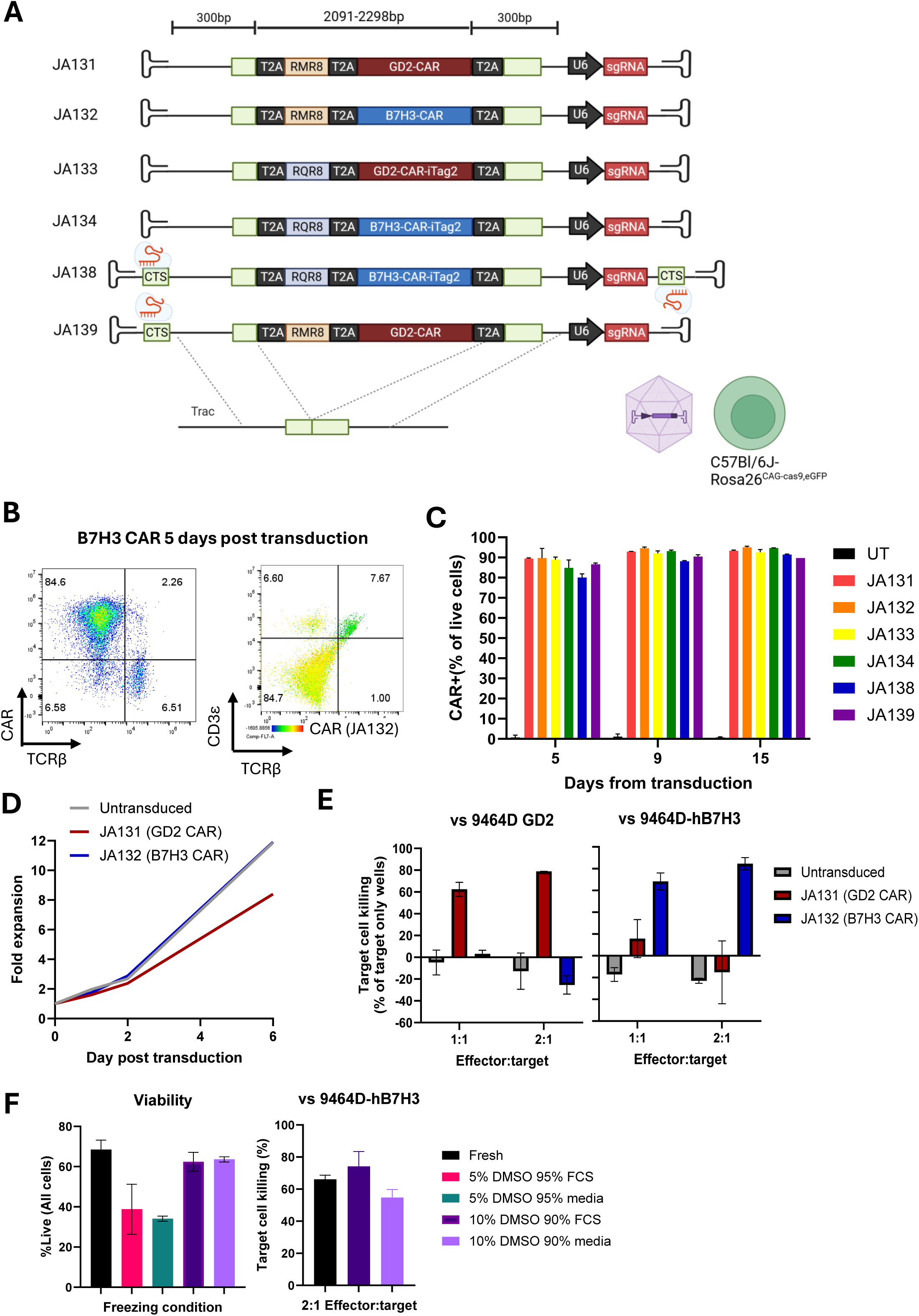
Cas9 mouse T cells were isolated and activated for 48 hours with soluble anti-CD28 and platebound anti-CD3. Cells were then transduced with AAV at MOI 1E5 in the presence of 1uM AZD7468 and 3uM POlQi2 for 24 hours before expansion in fresh media with IL2 IL7 and IL15. A) Schematic of CAR HDRT constructs used. All constructs had the same 300bp homology arms to the mouse Trac locus, but differed in their marker genes, CAR constructs and the presence of flanking Cas9 target sites (CTS). The marker genes RQR8 and RMR8 and the CARs targeting GD2 or human B7H3 with or without the degron tag, iTag2, are as described in the materials and methods. B) Representative flow cytometry 5 days post AAV transduction. Colour axis on right hand plot represents the degree of CAR positivity. Cells positive for CD3ε and CAR are presumed to be γδ-T cells. C) CAR expression of all constructs after extended in vitro culture. Data are mean of 2 technical replicates. UT = untransduced D) Expansion post transduction. Data are from 3 pooled spleens transduced at an MOI 1x10^5 with JA131 or JA132. E) 24 hour co-culture of GD2 CAR (JA131) or B7H3 CAR (JA132) with 9464D expressing either GD2 or human B7H3. Data mean of are 2 independent replicates performed in technical triplicate. Target cell killing was calculated relative to the mean cell number in target only wells. F) Flow cytometry assessment of cell viability and cytotoxicity for JA132 transduced cells following cryopreservation in different cryopreservation media compared to the same cells kept in culture for less than 2 weeks. Cytotoxicity assay is as in E but normalised to target cells in wells with untransduced T cells. Data are 2 technical replicates.

### TRAC-CARs show long term in vivo persistence

Our B7H3 targeting CAR has an scFv with specificity for the human B7H3 isoform. We therefore assessed the *in vivo* persistence of CAR T generated using AAV in an immune competent model using subcutaneous tumours that express human B7H3 under the transcriptional control of the murine B7H3 locus. Cells were engrafted in 129SvJ:C57Bl6-Rosa26eGFPCas9 mice to account for the mixed background of the 370566 tumour, and the need for a host tolerised to Cas9 to minimise MHC-dependent rejection (Figure 4A). Tumour bearing mice were administered intraperitoneal cyclophosphamide at a dose that leads to both lymphodepletion and tumour regression (Figure 4B and 4C). As the lymphodepletion alone caused substantial regression of the tumor, this model system is not yet able to evaluate the in vivo anti-tumor efficacy of these CAR-T cells. However it was possible to evaluate CAR-T persistence. 3 days post administration, CAR T cells represented a mean of 3% of total circulating leukocytes and as expected decreased (when measured as a % of CD45^+^ cells) after the monocytic rebound recovery from lymphodepletion on days 10 and 17 as monocytic lineages are more depleted by cyclophosphamide (Figure 4C and 4D). To assess the persistence of CAR T cells within the lymphocyte population, Thy1.2 was used as an alternate T cell marker (as the CARs lack CD3 or TCR expression). The proportion of T cells which were CAR positive remained stable up to 17 days post administration despite the expected tumour regression (Figure 4D).

**Figure 4.**
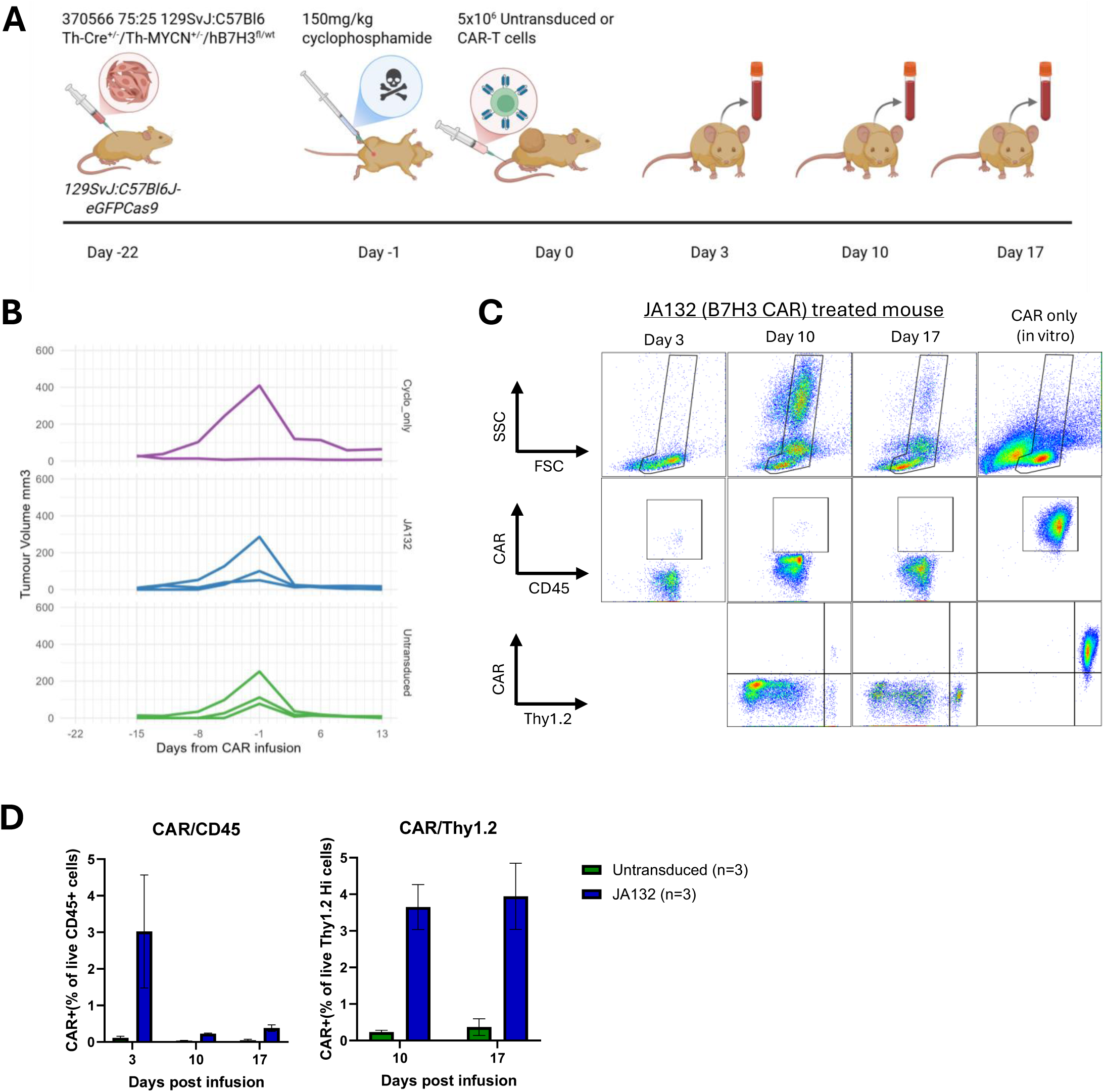
A) Experimental schematic: 370566 Neurospheres (75:25 129SvJ:C57Bl6 Th-Cre^+/-^/Th-MycN^+/-^/hB7H3 ^fl/wt^) were engrafted in the flank of female 129SvJ:C57BL6J-eGFPCas9 F1 mice. After 21 days all mice had palpable tumours and were administered 150mg/kg cyclophosphamide by intraperitoneal injection. 24 hours later 5x10^6 Untransduced or B7H3 CAR-T cells were administered by tail vein injection B) Tumour volume by calliper measurements. n =2 mice for cyclophosphamide only, n=3 mice for cyclophosphamide + JA132 (B7H3 CAR) and cyclophosphamide + untransduced T cells. C) Representative flow cytometry analysis of peripheral venous blood samples from lymphodepleted mice treated with JA 132. Top panel demonstrating predominant loss of granulocytes and monocytes 3 days after lymphodepletion with cyclophosphamide and subsequent recovery by day 10. Middle and bottom panels assess CAR percentage relative to total CD45 positive cells and to Thy1.2 high (T cells). Gating is based on FMOs and staining of in vitro cultured CARs (right most column) D) Persistence of CAR T cells in peripheral blood as a proportion of leukocytes (CD45 positive) or T cells (Thy1.2 high) n = 3 mice

## Discussion

CRISPR/Cas9 KI is an attractive method for development of CAR-T in immune competent mouse models. However there are differences between mouse and human T cells’ ability to be cultured and genetically manipulated ex vivo. dsDNA electroporation in human T cells negatively impacts cell viability, which can be partially ameliorated by inhibition of cGAS STING and use of proprietary HDR enhancers^29^. We confirm the observations of other groups that mouse T cells are more sensitive to dsDNA electroporation, necessitating the use of AAV as a HDRT. Use of the mouse T cell specific capsid Ark313 has previously been shown to significantly improve editing efficiency. We have identified further refinements to this approach. First, we identified that activation with plate-bound anti-CD3 and soluble anti-CD28 antibodies results in superior editing efficiencies than using anti-CD3/CD28 magnetic beads. Second, we identify the “goldilocks zone” for transduction post activation is 36-48 hours, the same as in human T-cells. Third we demonstrate that, in mouse T cells, the transient use of inhibitors of alternate mechanisms of DSB repair enhances editing efficiencies whilst maintaining good cell viability and expansion potential.

Given the space constraints of AAV (4.7kb including ITRs) using miniature promoters for guide RNA expression would increase the potential size of the HDRTs. This is particularly important for multi-component “armoured” CAR T cells. Unfortunately, we found the 96bp M11 promoter was less efficient than the 249bp U6 promoter. A miniaturised version of the U6 promoter which is 112bp has been shown to have comparable efficiency in a lentiviral construct^30^ and is worthy of evaluation in future work. Alternatively, targeting a HDR to the CD3z locus reduces the size of the CAR construct (as the CD3z endodomain is not required)^28^, increasing the side of possible armouring molecules/receptors.

Importantly, we have identified a cryopreservation protocol that results in retained cytotoxic functionality upon thawing. This will facilitate collaborative efforts as T cells can be generated at one centre and tested at centres with specific *in vivo* models and expertise.

Whilst the use of transgenic cas9 mice removes the need for RNP electroporation, which can reduce cell viability, a potential limitation for *in vivo* experiments is limited persistence due to MHC presentation of Cas9 protein triggering an immune response. However, adoptive transfer of CAR-T within the immune competent setting requires systemic lymphodepletion which is likely to prevent significant rejection in typical short term experiments. Moreover, we have mitigated rejection by using allograft hosts tolerised to Cas9. This can either be Cas9 mice for tumour cell lines from a pure C57Bl/6J background, or using F1 hybrids of C57Bl6-Cas9 and another strain for allografts on different backgrounds. An additional consideration when using hybrid mouse strains is the potential for graft versus host disease (GvHD) when administering T cells from C57Bl6-Cas9 mice into hybrid mice which are heterozygous for mismatched MHC alleles and may express allogenic antigens. This was not a concern in this study as both 129SvJ and C57Bl6 share the same MHC alleles, and the high knock-in efficiency meant that there were few TCR positive cells infused. Bead-based depletion of remaining positive TCR positive cells prior to infusion should both remove the possibility of GvHD and enrich the CAR population.

We have demonstrated persistence of administered T cells for over 2 weeks despite recovery from transient lymphodepletion. Because the tumor we used in these in vivo studies showed a strong response to the cyclophosphamide lympohodepleting chemotherapy alone, we could not yet analyze the anti-tumor efficacy of the CAR-T cells generated and described in this report. Studies are underway to test these same CAR-T cells in mice bearing separate syngeneic tumors that do not respond so well to the lymphodepleting chemotherapy; this ongoing work will be reported separately.

In summary, we have developed an optimised method for CRISPR/Cas9 knock-in for mouse T cells. Whilst we have focussed on the TRAC locus, our approach is likely to be applicable to other loci including genomic safe harbours for constructs with inducible promoters. By generating long lived cells with high transduction efficiency our approach can be used to assess CAR-T cell libraries and pharmacokinetic studies of drugs that modulate CAR expression within the tumour microenviroment.

## Acknowledgements

UCL GeneTxNeuro Vector Core Facility for initial training in AAV manufacture. Evon Poon, Karen Barker and Barbara Martins (Chesler Group) for breeding and provision of 370566 tumour line.

## Funding

TJJ is funded by an NIHR Clinical Lecturer award. This work was funded by an Academy of Medical Sciences Clinical Lecturer starter grant (SGL030\1009). SMT and CH are funded by the Little Princess Trust (grants 575187 G22 21DC39 and CCLGA 2025 01 Anderson) CRUK Stand up to Cancer (su2c RT6188) and CRUK Cancer grand challenge (CGCATF-2023/100051). JA salary is funded by the NIHR Great Ormond Street Hospital Biomedical research Centre, with lab supplies and infrastructure from grants from RICC and Children with Cancer UK. Generation of human B7H3 tumour models was funded by Neuroblastoma-UK.

## CreDiT contributions

TJJ: Conceptualisation, Funding acquisition, Investigation, Methodology, visualisation, Writing – Original Draft Preparation

SMT: Investigation, Writing – Review & Editing

CH: Investigation, Writing – Review & Editing

HB: Investigation,

FA: Investigation

LKD: Resources, Writing – Review & Editing

AKE: Resources, Writing – Review & Editing

PMS: Resources, Writing – Review & Editing

LC: Resources, Writing – Review & Editing

JA: Supervision, Funding acquisition, Writing – Review & Editing

## Conflicts of Interest

JA hold patents in CAR-T technology development, including a pending patent for the TE9 anti-B7H3 binder. JA holds founders shares in Autolus.

