## Supplementary figures for "An optimised method for generation of murine CAR-T cells by CRISPR/Cas9"

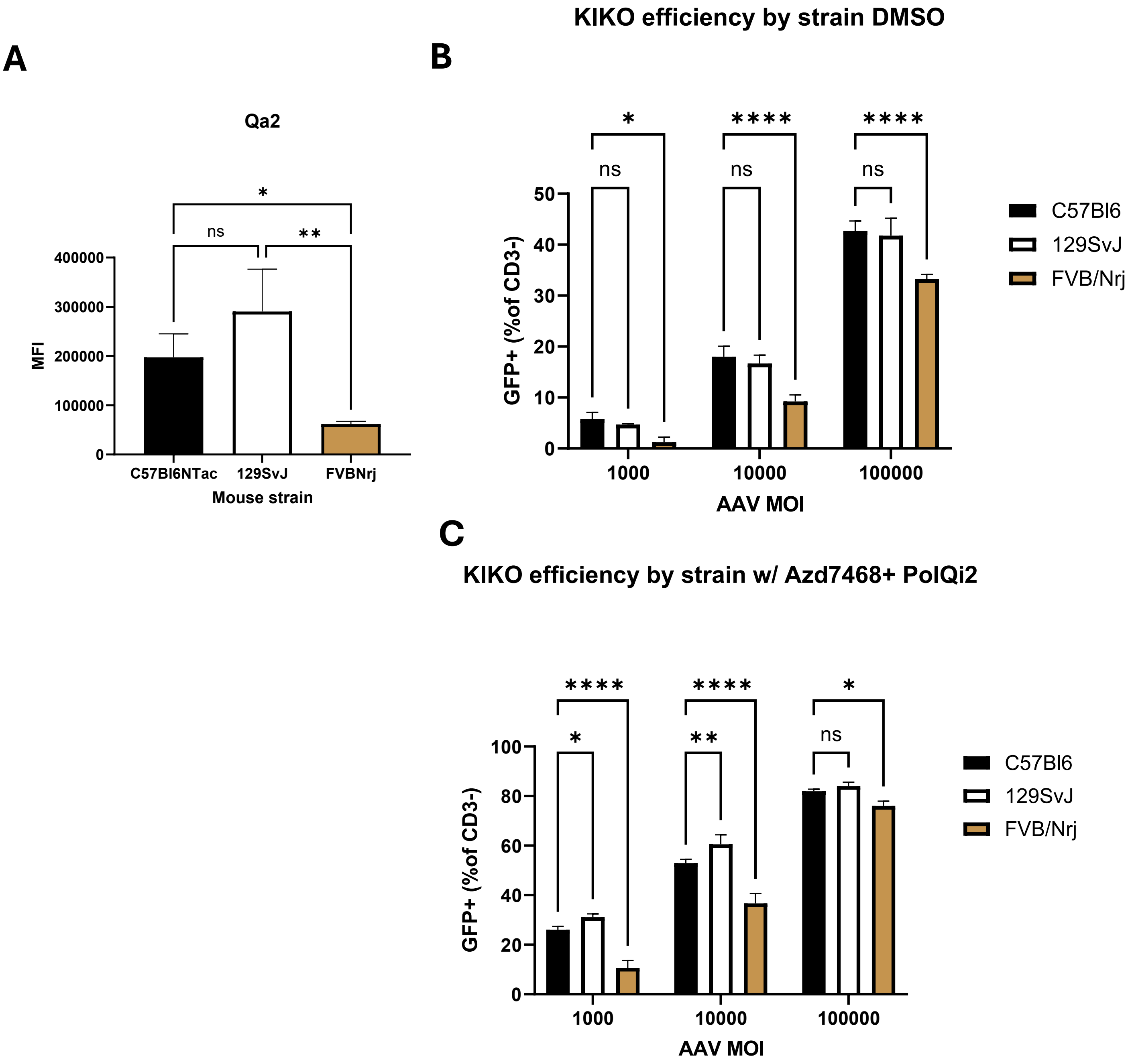


Supplementary Figure 1

A) Qa2 surface expression on three mouse strains quantified by median fluorescence intensity (MFI) of flow cytometry. Data are mean and standard errors of 3 technical replicates
B) Knock-in/Knock out efficiency using RNP electroporation and AAV HDRT by strain, Knock in (GFP positive) expressed as a proportion of knock-out (CD3 negative). Data are mean and standard errors of 3 technical replicates.
C) As for Figure B but AAV transduction performed in the presence of 1uM AZD7468 and 3uM PolQi2

A

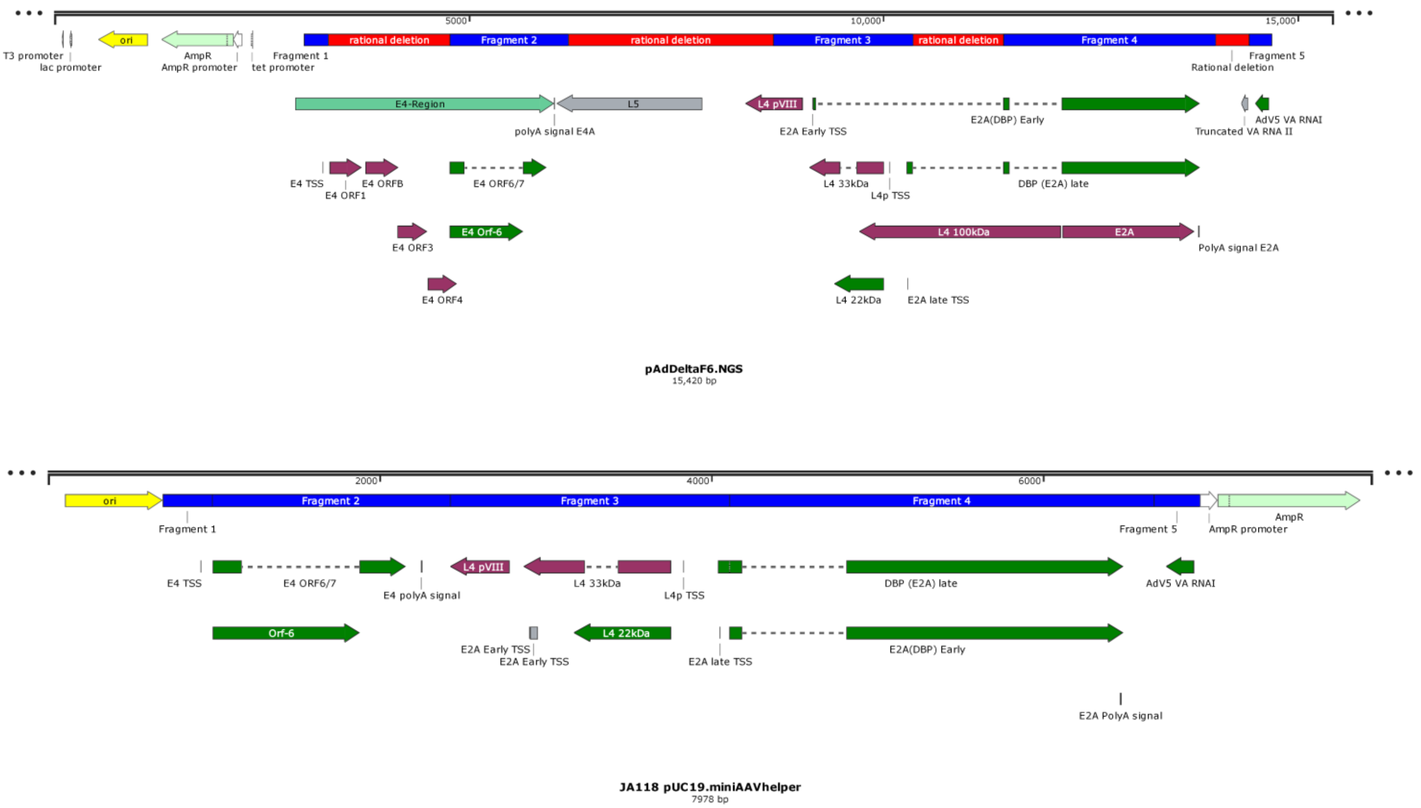


B

| Fragment | Contents | AdV5 Genomic location |
|---|---|---|
| 1 | E4 TSS | 35543-35834 |
| 2 | E4 ORF-6<br>E4 ORF-6/7 | 32646-34083 |
| 3 | E2A Early TSS<br>E2A late TSS<br>L4 TSS<br>L4-22kDa | 27528-25839 |
| 4 | E2A ORF | 24745- 22188 |
| 5 | VA RNA I | 10853-10570 |

C

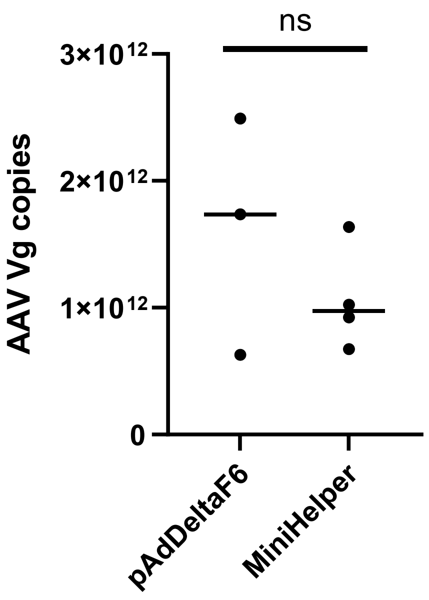


Supplementary Figure 2

A) Annotated linearised plasmid maps for pAdDeltaF6 and generated minihelper plasmid. Genes required for recombinant AAV production in HEK293Ts are highlighted in green

B) Location from fragments taken from pAdDeltaF6 relative to Adenovirus 5 genome (NCBI Reference Sequence: AC_000008)

C) AAV titre from small scale AAV preparations, 3 preps using pAdDeltaF6 and 4 using Minihelper. Titres were quantified by ITR qPCR and normalised to final prep volume to give absolute viral genome copy numbers.

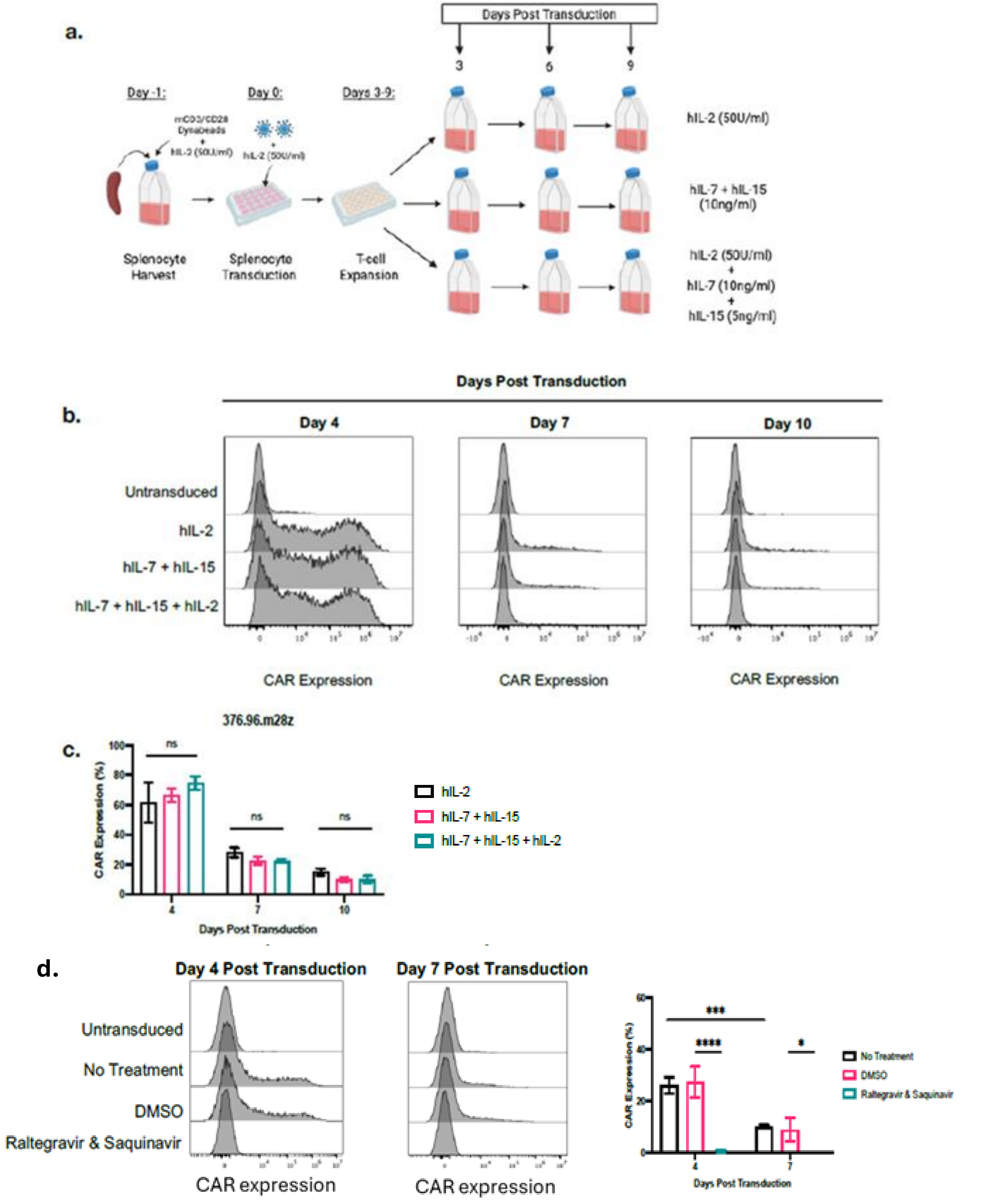


Supplementary Figure 3

A) Schematic of generation of fully murinised B7H3 targeting CAR (376.96.m28z) by retroviral transduction of 129SvJ splenocytes activated with mCD3/CD28 dynabeads. Retrovirus was generated using a Phoenix Eco producer line. Following transduction cells were expanded in either human IL2 only, human IL-7 and human IL-15 or all 3 cytokines.

B) Representative CAR expression by flow cytometry in different expansion media.

C) Longitudinal in vitro CAR expression from 3 independent mouse spleens, mean and standard error.

D) CAR expression after retroviral transduction in the presence or absence of retroviral protease inhibitor saquinavir and integrase inhibitor raltegravir. Data are presented as representative flow plots and mean and standard errors from 3 independent mouse spleens.

Supplementary table 1 – Flowcytometry antibodies used in this study

| Target | Clone | Flurochrome | Supplier | Catalog |
|---|---|---|---|---|
| **Mouse CD45** | 30-F11 | APC<br><br>BV510 | Biolegend | 103112<br><br>103137 |
| **Mouse CD3** | 17A2 | APC | Biolegend | 100236 |
| **Mouse TCRb** | H57-597 | BV711 | Biolegend | 109243 |
| **Human B7H3** | MIH42 | APC | Biolegend | 351005 |
| **GD2** | 14G2a | FITC | Biolegend | 357313 |
| **Thy1.2** | 30-H12 | APC | Biolegend | 105312 |
| **B7H3 CAR** | Recombinant B7H3-His tagged | | R&D biosystems | 1949-B3-050/CF |
| **Anti-His** | J095G46 | PE<br><br>APC | Biolegend | 362603<br><br>362605 |
| **GD2 CAR** | Protein L | PE | Biolegend | 303651 |
| **Qa2** | 695H1-9-9 | PE | Biolegend | 121715 |
